## Supplementary for "Thymic dendritic cell-derived IL-27p28 promotes the establishment of functional bias against IFN-γ production in newly generated CD4^+^ T cells through STAT1-related epigenetic mechanisms"

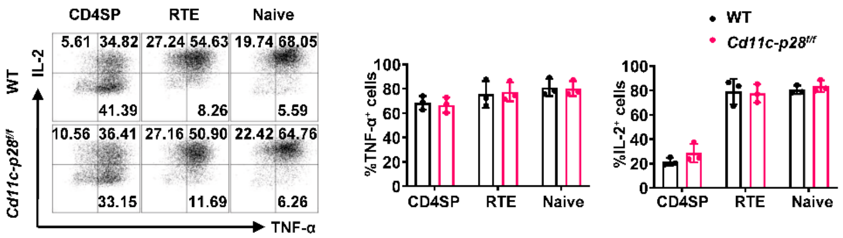


**Figure 1-figure supplement 1** The production of IL-2 and TNF-α was not altered during CD4SP thymocytes maturation for p28 deficiency.

CD4SP thymocytes, $CD4^+$ RTEs and naive $CD4^+$ T cells from *Cd11c-p28*$^{f/f}$ and WT mice were sorted, cultured under Th0 conditions for 3 days, and analyzed by intracellular staining. Representative dot plots (left) and statistical data (mean ± SD, right; n=3) were shown. No significant differences were observed (Student's *t*-test).

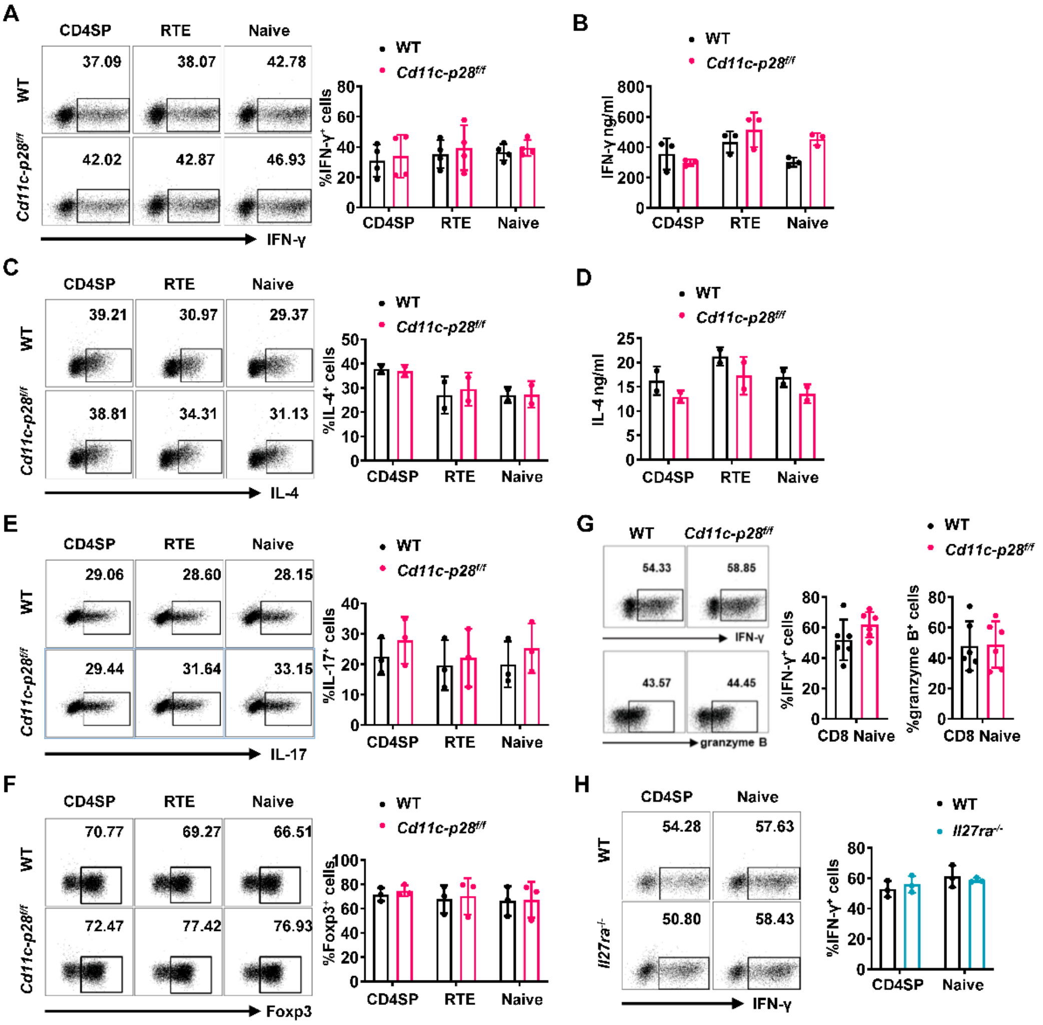


**Figure 1-figure supplement 2** *In vitro* differentiation of CD4$^+$ T cells under polarized conditions is unaffected by p28 deficiency.

Sorted CD4SP thymocytes, CD4$^+$ RTEs, and CD4$^+$ naive T cells from *Cd11c-p28*$^{f/f}$ and WT mice were cultured under Th1 (A-B), Th2 (C-D), Th17 (E) and Treg (F) conditions for 3 days.

(A) Frequency of IFN-γ$^+$ cells measured by intracellular staining. Representative dot plots (left) and statistical data (mean ± SD, right; n=4).

(B) IFN-γ concentration in the supernatants from Th1 cultures measured by ELISA (mean ± SD, n=3).

(C) Frequency of IL-4$^+$ cells measured by intracellular staining. Representative dot plots (left) and statistical data (mean ± SD, right; n=2).

(D) IL-4 concentration in supernatants from Th2 cultures measured by ELISA (mean ± SD, right; n=2).

(E-F) Frequency of IL-17A$^+$ (E) or Foxp3$^+$ (F) cells were measured by intracellular staining. Representative dot plots (left) and statistical data (mean ± SD, right; n=3).

(G) CD8+ naive T cells were cultured under Th0 conditions for 3 days. The frequency of IFN-γ-, and granzyme B-producing CD8$^+$ T cells were determined analyzed by intracellular staining. Representative dot plots (left) and quantification (right, mean ± SD, n=6).

(H) Frequency of IFN-γ$^+$ cells in CD4SP thymocytes and CD4$^+$ naive T cells from *Il27ra*$^{-/-}$ and WT mice cultured under Th1 conditions. Representative dot plots (top) and statistical data (mean ± SD, bottom; n=3). No significant differences were observed (Student’s *t* test).

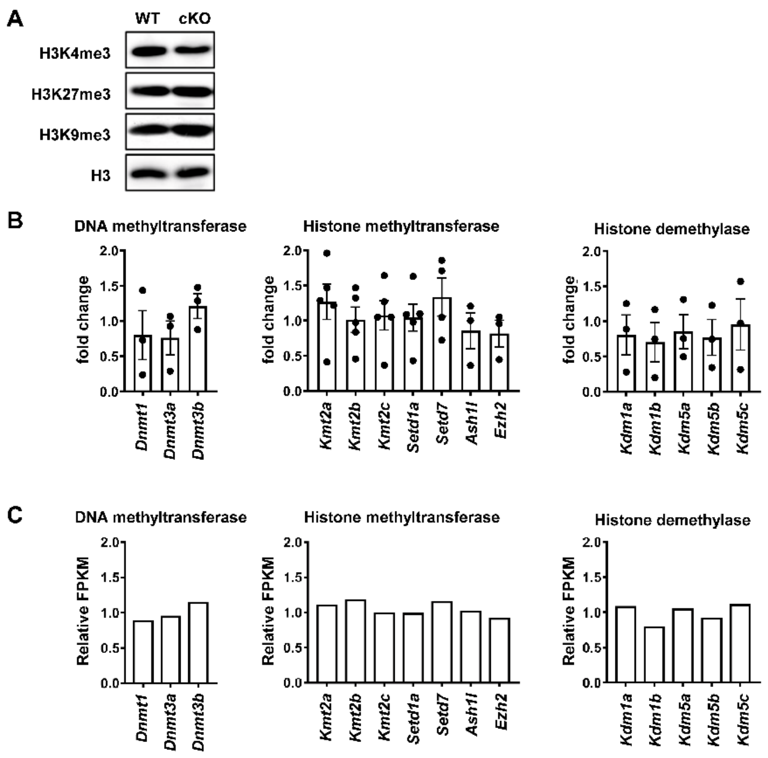


**Figure 3-figure supplement 1** Basal levels of H3K4me3, H3K27me3, H3K9me3, and methylation-related enzymes are unaffected by p28 deficiency.

(A) Global H3K4me3, H3K27me3, and H3K9me3 levels in $CD4^+$ naive T cells detected by Western blotting. H3 served as an internal control. Results are representative of three independent experiments.

(B) WT CD4SP thymocytes were sorted, stimulated with IL-27 (2 ng/mL) for 12 hours, and analyzed for mRNA levels of histone methyltransferases, demethylases, and DNA methyltransferases by qPCR. Data, shown as fold change over untreated cells (mean ± SEM; n=3-5), revealed no significant differences.

(C) Relative FPKM of epigenetic related enzymes in CD4SP thymocytes from RNA-seq data (Figure 4), shown as $FPKM_{Cd11c\text{-}p28f/f}/FPKM_{WT}$. No significant changes were observed.

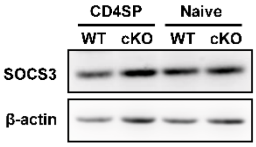


**Figure 5-figure supplement 1** SOCS3 levels in $CD4^+$ T cells from p28 deficient mice. CD4SP thymocytes and naive $CD4^+$ T cells were freshly isolated from WT and *Cd11c-p28*$^{f/f}$ mice, and SOCS3 expression was assessed by Western blotting. β-actin served as an internal control. Data are representative of three independent experiments.

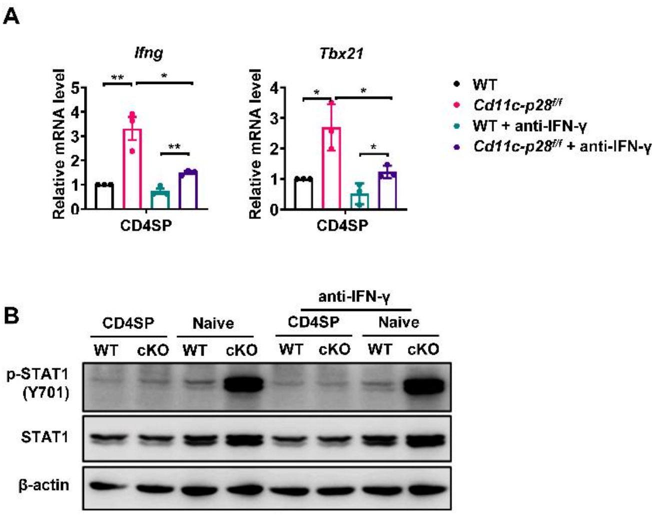


**Figure 5-figure supplement 2** The Stat1-dependent hyper-transcription of *Ifng* and *Tbx21* in p28 deficient CD4$^+$ T cells is not rescued by IFN-γ blockade.

(A) The CD4SP thymocytes and naive CD4$^+$ T cells from *Cd11c-p28*$^{f/f}$ and WT mice were stimulated with plate-bound anti-CD3 (2 μg/mL) and soluble anti-CD28 (1 μg/mL) without or with anti-IFN-γ (10 μg/mL) for 12 hours. *Ifng* and *Tbx21* mRNA levels were determined by qPCR. Data (mean ± SEM) are representative of three independent experiments. *, $p < 0.05$ and **, $p < 0.01$ (Student's *t*-test).

(B) The CD4SP thymocytes and CD4$^+$ naive T cells from *Cd11c-p28*$^{f/f}$ and WT mice were cultured without or with anti-IFN-γ (10 μg/mL) for 2 hours. STAT1 phosphorylation (pY701) was examined by Western blotting. Data are representative of three independent experiments.

**Supplementary Table 1** Primers used in this study.

| Gene | Forward Primer | Reverse Primer |
|---|---|---|
| *b-actin* | TATGGAATCCTGTGGCATC | GTGTTGGCATAGAGGTCTT |
| *Ifng* | TCAAGTGGCATAGATGTGGAAGAA | TGGCTCTGCAGGATTTTCATG |
| *Il4* | ACAGGAGAAGGGACGCCAT | GAAGCCCTACAGACGAGCTCA |
| *Il2* | TCCTGAGCAGGATGGAGAAT | GTCAAATCCAGAACATGCCG |
| *Tbx21* | TCAACCAGCACCAGACAGAG | ATCCTGTAATGGCTTGTGGG |
| *Gata3* | GCCTGCGGACTCTACCATAA | AGGATGTCCCTGCTCTCCTT |
| *Gm12250* | TAATGCCCTTCGGGGAATAGG | CTGGTTTGAAGTTAGTTGTCCCA |
| *Oasl2* | TTGTGCGGAGGATCAGGTACT | TGATGGTGTCGCAGTCTTTGA |
| *Usp18* | TTGGGCTCCTGAGGAAACC | CGATGTTGTGTAAACCAACCAGA |
| *Oas2* | TTGAAGAGGAATACATGCGGAAG | GGGTCTGCATTACTGGCACTT |
| *Oas3* | TCTGGGGTCGCTAAACATCAC | GATGACGAGTTCGACATCGGT |
| *Parp14* | AAGCAGATTGAAGTTGAGGACAA | CTTTGCCGGGGTTTCTGAAGT |
| *Ifit3* | TCAGGCTTACGTTGACAAGGT | CACACTTTAGGCGTGTCCATC |
| *Igtp* | CTCATCAGCCCGTGGTCTAAA | CACCGCCTTACCAATATCTTCAA |
| *Irf1* | ATGCCAATCACTCGAATGCG | TTGTATCGGCCTGTGTGAATG |
| *Ifi44* | AACTGACTGCTCGCAATAATGT | GTAACACAGCAATGCCTCTTGT |
| *Rsad2* | TGCTGGCTGAGAATAGCATTAGG | GCTGAGTGCTGTTCCCATCT |
| *Il12rb1* | ATGGCTGCTGCGTTGAGAA | AGCACTCATAGTCTGTCTTGGA |
